# Estrogen receptor alpha mediated repression of PRICKLE1 destabilizes REST and promotes uterine fibroid pathogenesis

**DOI:** 10.1101/2024.09.09.612036

**Authors:** Michelle M. McWilliams, Faezeh Koohestani, Wendy N. Jefferson, Sumedha Gunewardena, Kavya Shivashankar, Riley A. Wertenberger, Carmen J. Williams, T. Rajendra Kumar, Vargheese M. Chennathukuzhi

## Abstract

Uterine fibroids (leiomyomas), benign tumors of the myometrial smooth muscle layer, are present in over 75% of women, often causing severe pain, menorrhagia and reproductive dysfunction. The molecular pathogenesis of fibroids is poorly understood. We previously showed that the loss of REST (<u>RE</u>-1 <u>S</u>ilencing <u>T</u>ranscription factor), a tumor suppressor, in fibroids leads to activation of PI3K/AKT-mTOR pathway. We report here a critical link between estrogen receptor alpha (ERα) and the loss of REST, *via* PRICKLE1. PRICKLE1 expression is markedly lower in leiomyomas, and the suppression of PRICKLE1 significantly down regulates REST protein levels. Conversely, overexpression of PRICKLE1 resulted in the restoration of REST in cultured primary leiomyoma smooth muscle cells (LSMCs). Crucially, mice exposed neonatally to environmental estrogens, proven risk factors for fibroids, expressed lower levels of PRICKLE1 and REST in the myometrium. Using mice that lack either endogenous estrogen (*Lhb^-/-^* mice) or ERα (*Esr1^-/-^* mice), we demonstrate that *Prickle1* expression in the myometrium is suppressed by estrogen through ERα. Enhancer of zeste homolog 2 (EZH2) is known to participate in the repression of specific ERα target genes. Uterine leiomyomas express increased levels of EZH2 that inversely correlate with the expression of PRICKLE1. Using chromatin immunoprecipitation, we provide evidence for association of EZH2 with the *PRICKLE1* promoter and for hypermethylation of H3K27 within the regulatory region of *PRICKLE1* in leiomyomas. Additionally, siRNA mediated knockdown of EZH2 leads to restoration of PRICKLE1 in LSMCs. Collectively, our results identify a novel link between estrogen exposure and PRICKLE1/REST-regulated tumorigenic pathways in leiomyomas.

## INTRODUCTION

Uterine fibroids, also known as uterine leiomyomas (UL), are the most common tumors of the female reproductive tract. Uterine fibroids result from aberrant clonal expansion of smooth muscle cells in the myometrium and are a major health concern among women. It is estimated that the cumulative incidence of tumors by age 50 was greater than 80% for black women and nearly 70% for white women and over 25% of all women have clinical symptoms of pain, pressure, excessive bleeding, anemia and in some cases, infertility ^1–4^. Currently, the leading treatment for leiomyomas is hysterectomy, which is not only invasive and risky, but also presents a substantial financial burden. Leiomyomas accounted for up to $34.4 billion in medical costs in 2010 and continue to be the leading cause for hysterectomies in the United States ^5^. Despite the widespread need for a more efficacious treatment for leiomyoma, there is currently no approved pharmacotherapy treatment option for fibroids, which is long term, cost effective and that leaves fertility intact. Remarkably, the molecular mechanisms that trigger the pathogenesis of uterine leiomyoma are poorly understood.

Evidence from our laboratory and others indicates the central role of the PI3K/AKT/mTOR signal transduction pathway leading to cell growth, proliferation and cell survival in leiomyoma, as well as in malignant tumors ^6–8^. We reported previously that the loss of REST protein in uterine leiomyoma allows aberrant expression of GPR10 (PRLHR), a neuronal specific G-protein coupled receptor, leading to cell growth and survival via the PI3K/AKT-mTOR signal transduction pathway ^9^. Normally, GPR10 is silenced in non-neuronal tissues, including the myometrium, by its transcriptional silencer REST/NRSF (Repressor Element Silencing Transcription factor/ Neuron-Restrictive Silencing Factor) ^10,11^. REST is a major tumor suppressor that functions by epigenetic regulation of a multitude of its target genes through the recruitment of chromatin modifying enzymes to gene promoters ^11,12^. REST prevents the expression of neuron-specific genes in non-neuronal tissues and suppresses the PI3K/AKT pathway ^13^. In leiomyomas, the loss of REST leads to the critical activation of the PI3K/AKT-mTOR pathway, mediated by the de-repression of GPR10 and possibly by other REST target genes ^9^. In addition to GPR10, many of the most aberrantly expressed genes in leiomyomas are direct targets of REST, indicating that the loss of REST has an important role in downstream epigenetic changes leading to leiomyoma development and growth. Recently, we showed that conditional deletion of *Rest* in mouse uterus leads to UL phenotype, including tumor development, gene expression analogous to human UL, and revealed a direct role for REST in aberrant estrogen and progesterone signaling ^14^. The widespread expression of GPR10/ PRLHR in UL, demonstrated previously by us and others ^15^, imply near ubiquitous loss of REST in UL. Despite this, molecular mechanisms that promote the loss of REST protein in UL are unknown.

Pre-pubertal exposure to environmental estrogens including DES, bisphenol A (BPA) and genistein has been implicated as a primary risk factor for the development of uterine fibroids later in the reproductive life of women ^16–21^.

Studies in rodent models have indicated that long-lasting epigenetic modifications that accompany early exposure to estrogenic compounds may promote UL formation ^20,22^. Because REST is a major epigenetic regulator of long-term gene repression in the periphery, we hypothesized that environmental estrogens may promote the development of leiomyomas by lowering REST levels and thereby impairing its function in the myometrium. Interestingly, in leiomyomas the *REST* mRNA level is unchanged in comparison to healthy myometrium, indicating that REST expression is regulated post-transcriptionally. In differentiating neuronal progenitor cells, degradation of REST is mediated by the E3 ubiquitin ligase, β-TRCP, which targets it for proteasomal degradation ^23–25^. In addition, the loss of REST by the ubiquitin-proteasome pathway has been reported in various cancers ^13,26^. Intriguingly, despite normal β-TRCP levels in both myometrial smooth muscle cells (MSMCs) and leiomyoma smooth muscle cells (LSMCs), REST is degraded at a faster rate in cultured leiomyoma cells compared to normal myometrial cells ^9^.

Given the widespread loss of REST and the relatively normal levels of β- TRCP expression in leiomyomas, we hypothesized that alternative mechanisms exist for the functional loss of REST in leiomyoma cells. In an effort to identify the mechanism of loss of REST in leiomyoma, we focused on PRICKLE1, a WNT/PCP protein that associates with REST. We found that uterine leiomyomas expressed significantly lower levels of PRICKLE1, and its expression mirrored that of REST. PRICKLE1, also known as RILP (REST-interacting LIM domain protein), regulates nuclear localization of REST ^27,28^. Interestingly, the *Prickle1* promoter contains estrogen response elements (ERE) that are directly bound by ERα in the mouse uterus ^29^, presenting a potential link to the well-recognized role of environmental estrogens in the pathogenesis of uterine leiomyomas. Using a series of *in vitro* and *in vivo* methods, we demonstrate here that *Prickle1* is downregulated by environmental estrogens in the myometrium and that the loss of PRICKLE1 leads to the loss of REST in LSMCs. Additionally, we identify a direct role for the polycomb repressive complex protein EZH2 (enhancer of zeste homolog 2) in the repression of *PRICKLE1* in leiomyoma smooth muscle cells. Taken together, we identify a novel pathway that connects environmental estrogen exposure, suppression of PRICKLE1, loss of REST-dependent epigenetic control, and aberrant gene expression in leiomyomas.

## RESULTS

### PRICKLE1 is aberrantly expressed in fibroids in association with REST and REST target genes

To understand the molecular basis for the loss of REST in uterine leiomyomas, we undertook a candidate gene approach and investigated the status of REST-associated genes in human leiomyoma using gene expression profiling data available from the GEO dataset (GSE13319) ^6^. This dataset represents one of the most extensive gene expression studies available for uterine fibroids. Ingenuity™ Pathway Analysis (IPA) showed that many of the most significantly overexpressed genes in leiomyoma, such as GRIA2, GRIN2A, DCX, STMN2, NEFH, GPR10, SCG2 and SALL1 are all direct targets of REST-mediated long-term repression (Fig. 1A). Conversely, PRICKLE1 and HB-EGF, two of the REST pathway associated genes with putative roles in the regulation of REST, were down regulated in uterine leiomyomas (Fig. 1A). Gene expression analysis in replicates of matched myometrial and leiomyoma samples from patients confirmed the reduced expression of PRICKLE1 in ULs compared to that in myometrial tissue (Fig. 1B, Fig. S1). PRICKLE1 protein levels were also significantly reduced in leiomyoma as indicated by immunoblot (Fig. 1C, D) and immunofluorescence (Fig. 1E) analyses. Interestingly, reduced levels of PRICKLE1 also mirror the low REST levels in leiomyoma compared to myometrial tissue (Fig. 1C). Because PRICKLE1 associates directly with REST and is essential for REST function and nuclear localization ^27^, we explored the role of PRICKLE1 as a candidate for the regulation of REST stability, function and for the pathogenesis of fibroids. Importantly, while normal myometrial smooth muscle tissue sections show robust expression and nuclear localization of REST, the traces of REST that persist in leiomyoma samples have a predominantly cytoplasmic localization (Fig. S2 A, B), suggesting that PRICKLE1-mediated nuclear localization of REST may be dysfunctional in uterine fibroids. Furthermore, ectopically expressed REST-GFP fusion protein was unstable ^9^ and mislocalized to the cytoplasm in LSMCs instead of its normal nuclear localization observed in MSMCs (Fig. S2 C, D). These findings suggested that the reduced PRICKLE1 expression occurring in leiomyoma tumor cells may play a role in REST localization as well as stability. The combined loss of both PRICKLE1 and REST in leiomyomas and the mislocalization of REST in leiomyoma cells point to a direct role for PRICKLE1 in the loss of REST in uterine fibroids.

**Fig. 1.**
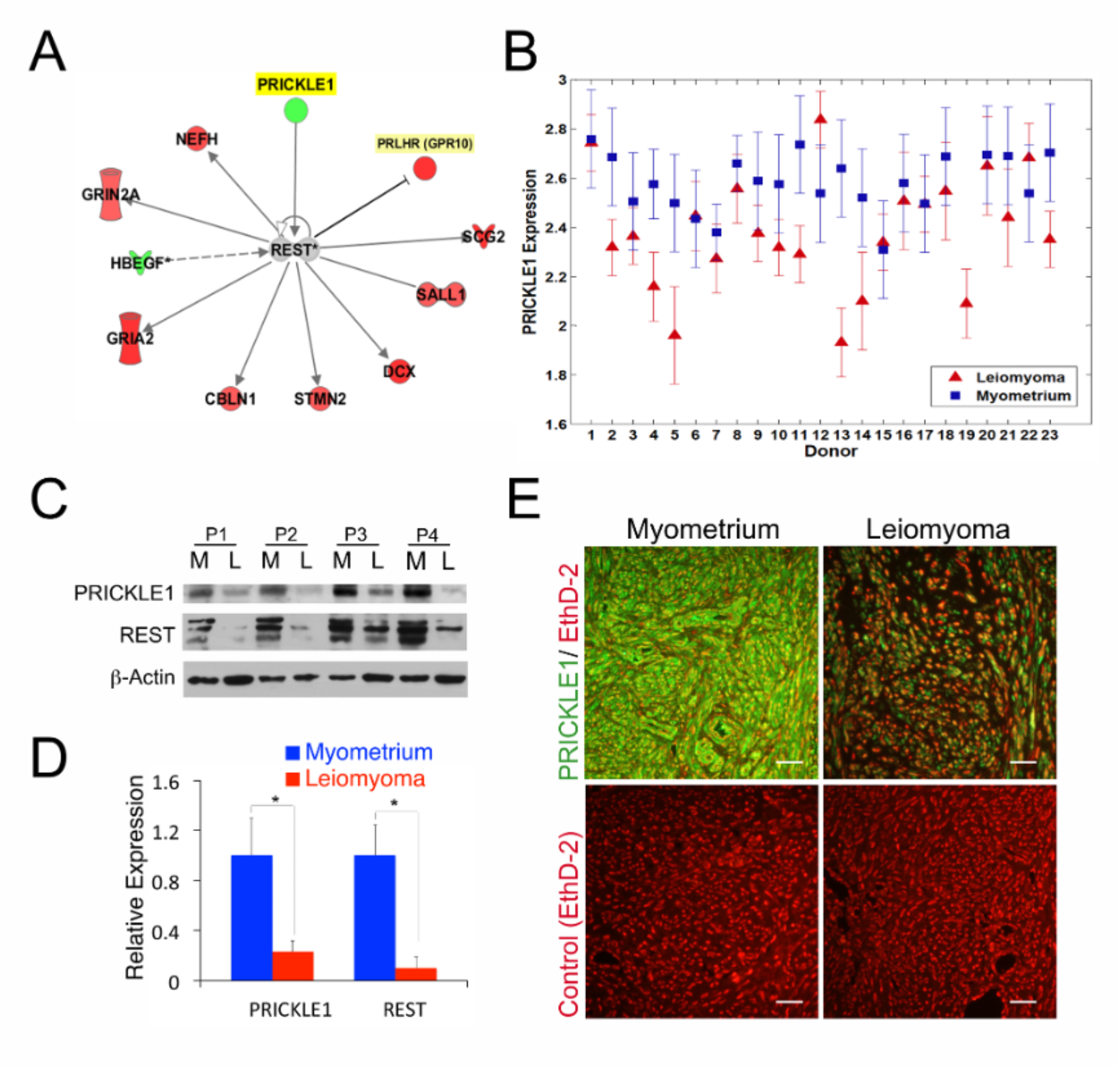
PRICKLE1, in association with REST, and REST-target genes are altered in uterine leiomyomas. (A) Ingenuity™ Pathway Analysis of REST associated genes in uterine leiomyoma tissues using gene expression profiling dataset (GSE # 13319). (B) *PRICKLE1* gene expression in uterine leiomyoma and myometrial patient samples in GSE # 13319. Immunoblot analysis (C) and densitometry (D) of PRICKLE1 and REST expression in uterine leiomyoma and myometrial tissues. β-Actin was used as loading control. (E) Immunofluorescence analysis of PRICKLE1 (green) expression in myometrial and leiomyoma tissue samples. Nuclei were stained with EthD-2 (red). * *P* < 0.05. Magnification bars represent 50μm.

### PRICKLE1 regulates REST expression in uterine leiomyomas at protein level

To elucidate the role of PRICKLE1 in REST stability and to establish a molecular mechanism for their concomitant loss in uterine leiomyomas, we tested whether modulating PRICKLE1 expression levels had an effect on REST protein levels. We silenced *PRICKLE1* expression in cultured primary MSMCs using siRNA transfection and probed for REST expression (Fig. 2A). PRICKLE1 knockdown caused a corresponding decrease in REST protein levels. Furthermore, overexpression of FLAG- tagged PRICKLE1 in cultured primary LSMCs caused an increase in REST protein levels (Fig. 2B). To confirm that REST regulation by PRICKLE1 occurs at post-transcriptional level as inferred from patient data, we further analyzed *PRICKLE1* expression vector-transfected LSMCs by quantitative PCR (Fig. S3). As expected, *PRICKLE1* mRNA expression was significantly increased in transfected cells, but there was no change in *REST* mRNA, indicating that REST protein stability, but not its mRNA expression was influenced by PRICKLE1. These data suggest that PRICKLE1, in addition to its role in the nuclear localization of REST, functions to stabilize REST in the myometrium, and that loss of PRICKLE1 leads to the destabilization and loss of REST protein, without a corresponding down regulation of its mRNA in leiomyomas.

**Fig. 2.**
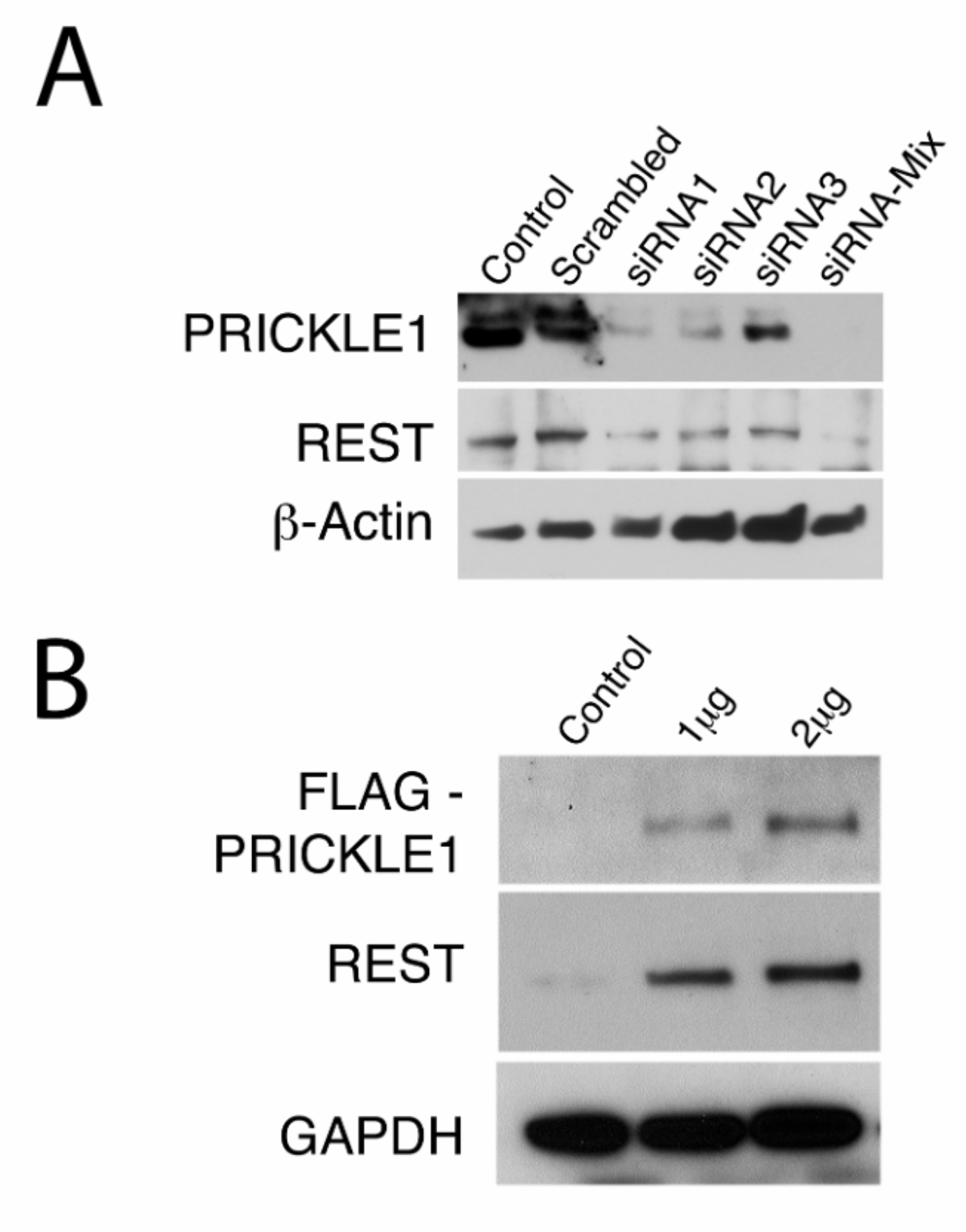
PRICKLE1 regulates REST expression in uterine leiomyomas. (A) Immunoblot analysis of REST expression upon *PRICKLE1* silencing in MSMCs. β-Actin was used as loading control. (B) Analysis of REST expression in LSMCs overexpressing FLAG-PRICKLE1. GAPDH was used as loading control.

### Estrogen receptor α suppresses PRICKLE1 expression in the myometrium

Pre-pubertal exposure to estrogenic chemicals has been proposed as a major risk factor for leiomyoma ^17,19,21,30^. In an effort to determine the sensitivity of PRICKLE1-REST pathway to estrogen exposure *in vivo*, we examined mice treated on postnatal days (PND) 1 - 5 with genistein, a common dietary phytoestrogen. Immunofluorescent staining for REST in PND5 uteri from genistein-treated and vehicle-treated control mice showed that mice exposed neonatally to genistein had significantly reduced levels of REST expression (Fig. 3A, B). Further, immunostaining for PRICKLE1 in these tissues showed markedly lower expression of PRICKLE1 in the myometrium (Fig. 3C, D), suggesting that early exposure to estrogenic chemicals may have significant effects on PRICKLE1 – REST mediated epigenetic control of gene expression in the uterus.

**Fig. 3.**
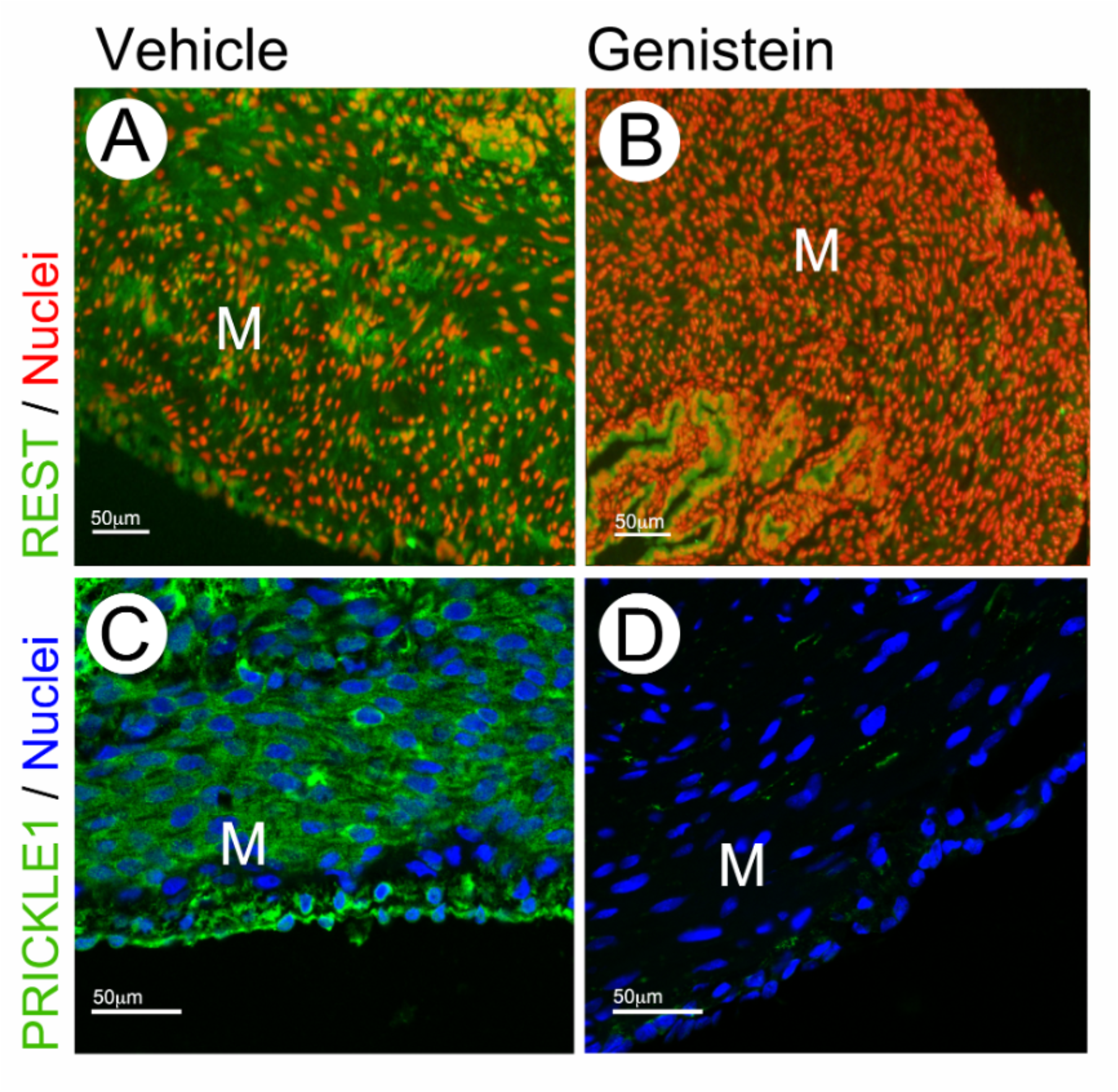
Phytoestrogen alters PRICKLE1 expression. Immunofluorescence analysis of PRICKLE1 (green) and REST (green) expression in vehicle (A &C) and genistein-treated (B & D) mouse myometrium. Nuclei were stained with EthD-2 (red) or DR (blue). Magnification bars represent 50μm. M: myometrium.

To test the putative direct role of estradiol in the regulation of PRICKLE1 expression in the mouse uterus we used luteinizing hormone (LH) beta-null (*Lhb^-/-^*), and hence LH-deficient, mice that have extremely low estrogen levels ^31^. PRICKLE1 expression in the myometrium of uteri from *Lhb^-/-^* mice at 12 weeks of age was significantly higher than in WT controls at estrus (Fig. 4A).

**Fig. 4.**
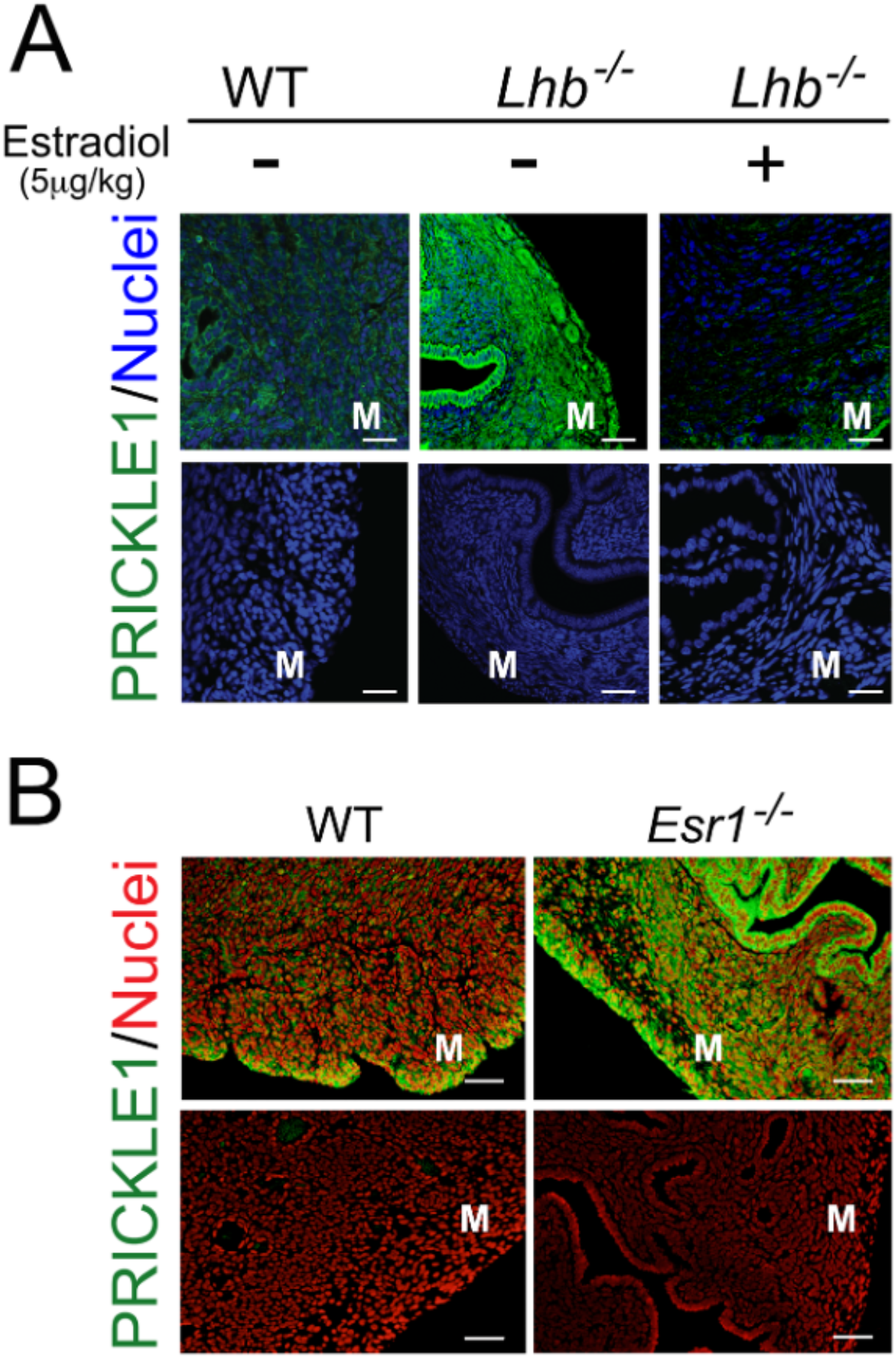
ERα regulates PRICKLE1 expression. (A) Immunofluorescence analysis of PRICKLE1 (green) in *Lhb* null mouse myometrium treated with or without estradiol. Nuclei were stained with DR (blue). (B) Expression of PRICKLE1 in WT and *Esr1* null mouse myometrium. Nuclei were stained with EthD-2 (red). Magnification bars represent 50μm. M: myometrium.

Importantly, uteri of *Lhb^-/-^* mice treated with estradiol (17-β estradiol, 5μg/kg, i.p. injected once daily for 5 days) reverted PRICKLE1 levels to the significantly reduced levels observed in wild-type mice (Fig. 4A), confirming a substantial down-regulation of PRICKLE1 expression by estrogen.

Data from an available ChIP-sequencing study showed that in the adult mouse uterus, estrogen response element (ERE) sequences proximal to the transcriptional start sites of *Prickle1* are associated with ERα ^29^. Thus, this association suggests that PRICKLE1 is a potential direct link between environmental estrogens and downstream tumorigenic pathways in the myometrium. To distinguish between the possible mechanisms of estrogen regulation of PRICKLE1, we examined *Esr1*-null mouse myometrium using fluorescent immunohistochemistry. In the absence of ERα, PRICKLE1 expression was significantly increased (Fig. 4B) despite very high circulating estrogen levels in these mice ^32^, indicating that ERα mediates estrogen-induced repression of PRICKLE1. In support of environmental estrogen-mediated suppression of *Prickle1* transcription, analysis of microarray data (GSE104402, GSE104401) from neonatal (PND5) mice treated with DES showed significant reduction in *Prickle1* mRNA (Fig. S4). In order to determine if neonatal exposure to DES had a lasting impact on the expression of *Prickle1*, mice treated with either vehicle or DES from PND 1 to 5 were ovariectomized at 8 weeks of age and 17β-estradiol or vehicle was administered 14 days after the surgery. TaqMan qRT-PCR assays comparing the expression of *Prickle1* in the uteri of vehicle or estrogen treated adult ovx mice confirmed the suppression of Prickle1 by estradiol (Fig. S5). Most importantly, even in the absence of 17β-estradiol treatment, mice treated neonatally with DES expressed significantly lower levels of Prickle1 compared to those treated with vehicle, suggesting a long-term impact of environmental estrogen exposure (Fig. S5). Suppression of *Prickle1* expression by 17β-estradiol in DES treated mice was most robust (over 80% at 24h) when compared to mice treated neonatally with vehicle (Fig. S5).

### Enhancer of zeste homolog 2 (EZH2) participates in the repression of PRICKLE1

In hepatocellular carcinoma cell lines, EZH2, the histone methyltransferase subunit of the polycomb repressor complex, binds to PRICKLE1 promoter and suppresses its expression ^33^. EZH2 also functions in the repression of ER target genes through association with prohibitin 2 (Phb2 or REA, repressor of estrogen receptor activity) an estrogen receptor co-repressor^34^. We analyzed the expression status and a potential role of EZH2 in the repression of PRICKLE1 in uterine leiomyomas. Analysis of GEO dataset GSE: 13319 showed that EZH2 mRNA levels were significantly increased in leiomyoma samples compared to those in normal myometrium, confirming an inverse correlation with PRICKLE1 expression (Fig. 5A). TaqMan qRT-PCR analysis of matched human myometrium and leiomyoma samples confirmed the increased expression of EZH2 (Fig. 5B). Further, western blotting analysis of patient samples showed that EZH2 protein expression was higher in leiomyoma samples compared to patient matched normal myometrium with a consistent inverse relation to PRICKLE1 expression (Fig. 5C, D). Furthermore, siRNA knockdown of EZH2 in cultured primary LSMCs resulted in the increase of PRICKLE1 expression (Fig. 5E), showing that EZH2 is overexpressed in leiomyomas, and may play a role in the repression of PRICKLE1.

**Fig. 5.**
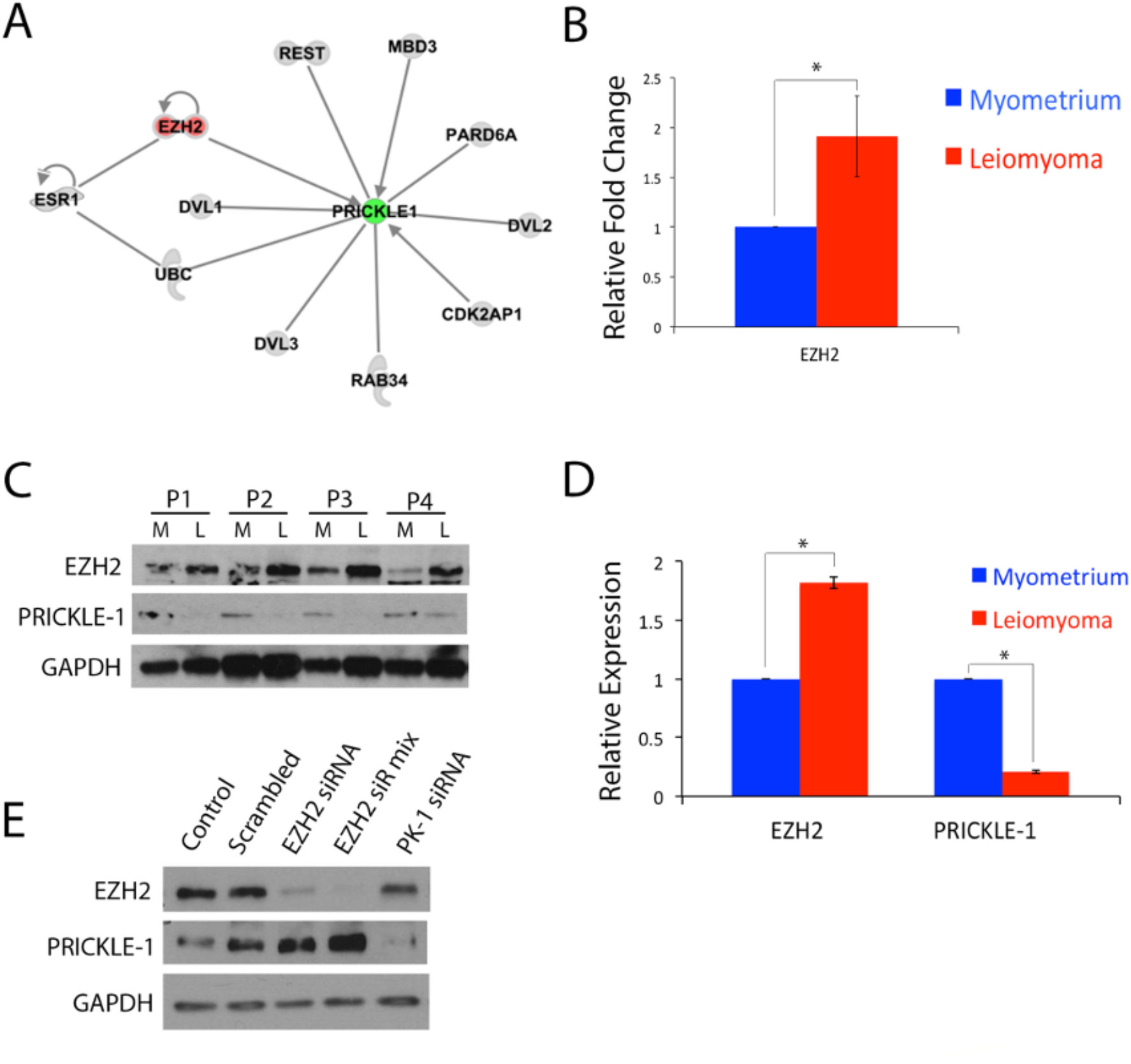
EZH2 mediates estrogen suppression of PRICKLE1 in leiomyomas. (A) Gene network analysis of *PRICKLE1* associated genes in uterine leiomyomas. (B) Gene expression analysis of *EZH2* in leiomyoma and myometrial tissues. (C) Immunoblot analysis of EZH2 and PRICKLE1 in matched human leiomyoma and myometrial tissues, and densitometry analysis (D) of western blot. (E) PRICKLE1 and EZH2 expression in EZH2-silenced LSMCs. GAPDH was used as loading control. * *P* < 0.05

We performed semi-quantitative ChIP-PCR experiments to identify a potential role of EZH2 mediated repressive H3K27 methylation in the regulation of PRICKLE1 expression in UL. Based on the UCSC genome browser data, PRICKLE1 is expressed with two major transcriptional start sites. The distal start site has overlapping ESR1 binding sites based on ENCODE factorbook motifs, as well as ReMap Atlas regulatory regions GSE132426.ESR1.Ishikawa_DMSO_D538G_clone3, GSE68355.ESR1.MCF-7_PROG and both the major transcriptional start sites are associated with tri- methylation of H3K27 in various cell lines (Fig. S6, indicated by red box).

Interestingly, the distal site was associated with increased H3K27Me3 and EZH2 in leiomyoma samples (Fig. S6 B, C), showing a putative role for EZH2 in the epigenetic regulation of *PRICKLE1* in UL. The proximal site (indicated by the blue box) did not show significant difference in H3K27Me3 between normal myometrium and Leiomyoma samples (data not shown).

### PRICKLE1 - WNT/ PCP pathway is dysregulated in uterine fibroids while canonical WNT/ β-catenin pathway remains unchanged

Since PRICKLE1 is an important mediator of WNT/PCP signaling and is reported to negatively regulate WNT/ β-catenin through ubiquitination of dissheveled 3 ^35^, we wanted to investigate if the loss of PRICKLE1 expression is related to the widely reported aberrant activation of WNT/ β-catenin pathway in uterine fibroids. Contrary to the belief that β-catenin activation promotes UL, we found no evidence for nuclear localization of β-catenin in UL (Fig. S7 A). Since patient to patient variation in protein expression or localization is possible, we repeated this experiment in over 50 patient samples and found no evidence for nuclear accumulation of β-catenin in UL (data not shown). We further investigated if TCF/LEF targets, which represent the activation of WNT/ β-catenin pathway are upregulated in UL from available GEO datasets (Fig. S7 B, GEO dataset GSE: 13319) and found no evidence for aberrant activation of canonical WNT/ β-catenin pathway. These results were confirmed using TaqMan qRT-PCR, where β-catenin and cyclin D1 were the only genes upregulated in this vast pathway. Since β-catenin mRNA is consistently upregulated in UL, we performed western blotting to investigate if LRP5/6 phosphorylation, which is indispensable in canonical WNT pathway activation, is dysregulated in UL. Our results indicated no evidence for hyper activation of LRP5/6 phosphorylation (Fig. S7 C). Interestingly, disheveled 1 (DVL1) mRNA and protein were significantly downregulated in UL samples while DVL2 and DVL3 remained unchanged (Fig. S7 C,E,F). Intriguingly, WNT inhibitory factors, SFRP1 and WIF1 were significantly upregulated in UL (Fig. S7 G). In contrast, several molecules involved in WNT/PCP, including WNT5A, FZD2, VANGL2, and MMP11 were significanly upregulated in UL (Fig S7. H). Additionally, there was increase in CDC42 phosphorylation in UL (Fig. S7 I).

## Discussion

The pathogenesis of uterine fibroids is poorly understood despite the widespread occurrence ^2^, significant morbidity ^1^, and the ever expanding medical cost ^5^ associated with the disease. While near ubiquitous existence of specific somatic missense DNA mutations ^36,37^ or chromosomal translocations ^38^ have been reported in uterine leiomyomas, the mechanisms that initiate those events in the relatively quiescent myometrial smooth muscle tissue have not yet been identified. We recently reported that the loss of tumor suppressor REST in uterine myometrial smooth muscle cells leads to aberrant gene expression and promotes the pathogenesis of uterine fibroids ^9^. REST, via direct binding to the RE-1 sequence elements in upstream regulatory regions, controls the long-term silencing of about 2000 genes by epigenetic mechanisms ^39,40^. The loss of REST by β−TrCP mediated ubiquitination and proteasomal degradation is suggested to result in PI3K/AKT dependent tumor cell proliferation and cell survival ^13,26^.

Intriguingly, the loss of REST in leiomyomas was not associated with enhanced β−TrCP mediated ubiquitin-proteasomal degradation of REST ^9^, suggesting that a novel mechanism may exist for its loss in leiomyomas.

We examined the regulation of known REST interacting proteins in leiomyomas in a candidate gene approach to identify the mechanism for the loss of REST in leiomyomas. Interestingly, PRICKLE1, an interacting partner of REST involved in its nuclear localization ^27^, was significantly down regulated in uterine leiomyomas (Fig. 1A, B). Further, REST protein showed predominant cytoplasmic localization in leiomyomas (Fig. 1C, Fig. S2B), suggesting that the loss of these protein partners may be linked. We had reported earlier that, in LSMCs transfected with a *GFP-REST* construct, the GFP-tagged REST protein was unstable and had failed to localize to the nucleus ^9^, indicating that the nuclear localization of REST was specifically affected in uterine fibroids. An inherited homozygous mutation on PRICKLE1 that disrupts its interaction with REST was reported to cause progressive myoclonus epilepsy (PME) with symptoms of neurological decline, including ataxia and dementia ^41^. PRICKLE1, also known as the REST interacting LIM domain protein (RILP), is a key regulator of the noncanonical WNT/PCP pathway ^42^. PRICKLE1 plays a crucial role during mouse embryo development by regulating the expression of VANGL2, BMP4, FGF8 and WNT5a ^42^. A function blocking mutation to PRICKLE1 (C251X/C251X) in mice results in major phenotypic changes in tissues of mesenchymal lineage ^42^. Most relevant to the well-recognized link between environmental estrogen exposure and the pathogenesis of uterine fibroids, a recent ChIP-sequencing study revealed that ERα is associated with the *Prickle1* promoter in the mouse uterus ^29^.

We hypothesized that the loss of PRICKLE1 expression in LSMCs is mechanistically linked to the loss of REST and smooth muscle tumor development in the uterus. We observed that upon siRNA-mediated knockdown of PRICKLE1 in cultured primary MSMCs, REST expression was correspondingly reduced (Fig. 2 A), indicating that the low PRICKLE1 levels in leiomyomas may indeed trigger the decline in REST stability and function.

Conversely, overexpression of Flag-tagged PRICKLE1 in cultured primary LSMCs led to a matching restoration of REST expression (Fig. 2B) without altering *REST* mRNA expression, confirming that the loss of PRICKLE1 expression in leiomyoma is linked to the destabilization of REST. Relevant to the role of PRICKLE1 in regulating the stability of REST without affecting its mRNA expression, we had shown earlier that the loss of REST protein expression in leiomyomas does not accompany a corresponding decline in its mRNA expression ^9^. Given the function of PRICKLE1 in the nuclear localization of REST, these results further support our observation that GFP-REST failed to localize to the nuclei of primary LSMCs upon transfection. The loss of normal REST function in various pathological states has thus far been linked to genetic mutations or to β−TrCP mediated degradation ^43–45^. Also, while the loss of REST function has been linked recently to the development of Alzheimer’s disease, a consistent mechanism for the loss of REST has not been identified ^46,47^.

Intriguingly, similar to the absence of its nuclear localization in leiomyoma, REST is lost from the nuclei of neurons in Alzheimer’s disease, instead appearing in autophagosomes along with pathological misfolded proteins. Further research will be needed to determine if the loss of PRICKLE1 leads to the degradation of REST in autophagosomes. The widespread destabilization and loss of REST in the absence of PRICKLE1 expression seen here in uterine leiomyomas is entirely novel, and may provide future therapeutic approaches for restoring REST function.

Somatic missense mutations to the mediator complex gene *MED12* have been reported in leiomyoma ^36^. MED12 is known to interact with REST through G9a, a methyl transferase responsible for repressive H3K9 mono- and di- methylation ^48^. Additionally, a *Med12* hypomorphic mutation in mouse embryos results in asymmetric distribution of PRICKLE1 and disruption of the WNT/PCP pathway ^49^. How the missense mutations to *MED12* reported in leiomyoma affect PRICKLE1 and the WNT/PCP pathway, in addition to REST localization and function are currently unknown.

Pre-pubertal exposure to environmental or dietary phytoestrogens such as DES, BPA, and genistein has been suggested to predispose women to the development of uterine fibroids ^18,19,21,30,50^. Also, the long-term effects of such endocrine disruptors in the uterus suggest the involvement of epigenetic gene regulatory mechanisms as has been shown in mouse models ^51^. Because REST is a major epigenetic regulator of gene expression, we investigated the impact of neonatal estrogen exposure on PRICKLE1-REST pathway. Our results indicated that the expression of PRICKLE1 and REST in the mouse uterus is negatively regulated by genistein (Fig. 3). Collectively, our data from the *Lhb* -/- and the *Esr1-/-* mouse models show that estrogen suppresses Prickle1 expression through ERα. Our data from mice treated neonatally with DES indicate long-term repression of *Prickle1* expression (Fig. S5) by environmental estrogens. Given the critical role that PRICKLE1 plays in the stability of REST (Fig. 2), the long-term repression of this tumor suppressor pathway by environmental estrogens could trigger genetic and epigenetic instability that promotes uterine fibroid pathogenesis.

Enhancer of zeste homolog 2 (EZH2), a polycomb group histone methyltransferase, regulates the expression of estrogen responsive genes in breast and prostate cancer cells via its association with the repressor of estrogen activity (REA, Prohibitin 2), an estrogen receptor co-repressor ^34^. Additionally, EZH2 directly binds to the promoter and negatively regulates the expression of PRICKLE1 in hepatocellular carcinoma cells ^33^, evoking the possibility for both EZH2 and ERα together playing a role in the suppression of PRICKLE1 in leiomyoma. Intriguingly, based on studies using ELT3 cells, it has been suggested that the down regulation of EZH2 by environmental estrogens via non-genomic pathways promotes the pathogenesis of uterine leiomyomas ^52^, though no patient data to indicate the suppression of EZH2 in leiomyoma was presented. In contrast to this hypothesis, our results indicated that EZH2 expression is significantly upregulated in leiomyoma samples both at mRNA and protein levels and this upregulation was inversely correlated with the expression of PRICKLE1 and REST (Fig. 5 B, C, D). Most importantly, the knockdown of EZH2 using siRNA resulted in the restoration of PRICKLE1 in leiomyoma cells (Fig. 5 E). Our results indicate that ERα - EZH2 mediated regulation of PRICKLE1 suppression in UL may involve changes in H3K27 methylation in its promoter. Our results support a novel ERα/ EZH2 – PRICKLE1 mediated, REST- target-specific gene derepression model (Fig. 6) in the pathogenesis of leiomyomas.

**Fig. 6.**
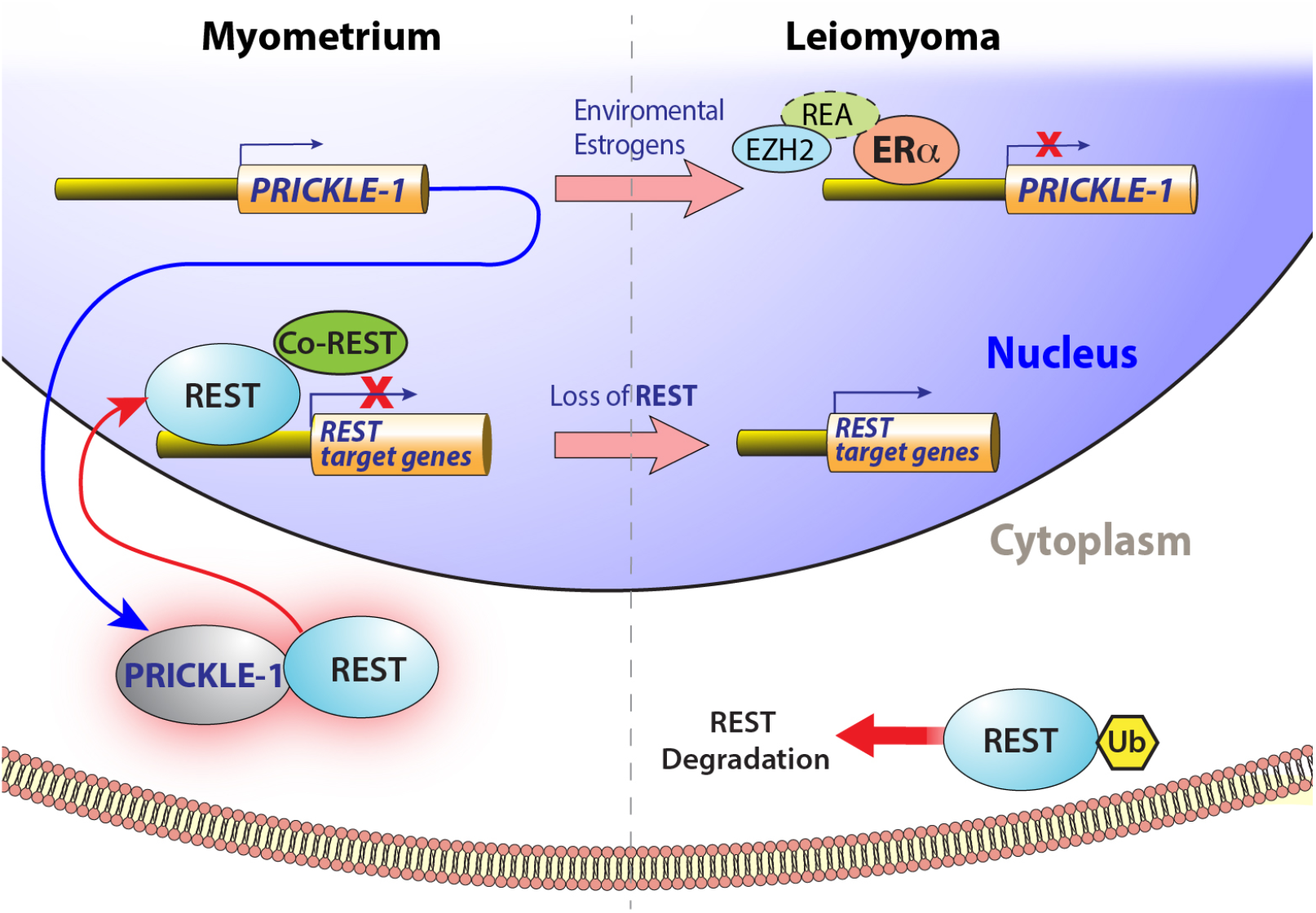
Regulation of PRICKLE1 and REST in leiomyomas. Schematic showing the regulation of *PRICKLE1* by ERα and regulation of REST by PRICKLE-1 in myometrium and leiomyoma.

There is a wealth of literature on the activation of WNT/β-catenin pathway in UL, including the activation in *MED12* mutant UL ^53^. Many of these studies are based on data showing upregulation of β-catenin mRNA in UL, which was also evident in our analysis (Fig. S7 B). Many, if not all, of the studies that include *in vitro* examination of WNT/ β-catenin pathway have utilized UL cell lines, which are highly unreliable, to investigate this pathway. Our data from UL tissue samples indicated no evidence for widespread activation of canonical WNT pathway in UL. Instead, WNT inhibitory molecules were found to be overexpressed in UL samples. Available gene expression datasets also indicated that TCF/LEF targets, downstream of WNT/ β-catenin were not overexpressed in UL (Fig S7. B). Increased cyclin D1 expression, observed in UL gene expression datasets (Fig. S7 B), may also arise independent of WNT/ β- catenin pathway ^54^. Similar to our results, scRNA data (GSE162122) from *MED12* mutant UL samples indicated no evidence for aberrant activation of WNT/ β-catenin pathway in any of the cell types present in tumors ^55^. Instead, WNT inhibitory molecules were upregulated in these tumor samples and negative regulation WNT/ β-catenin pathway was observed in *MED12* mutant UL associated fibroblast and smooth muscle cells ^55^. Additionally, suppression of canonical WNT/ β-catenin signaling, not activation of this pathway was reported previously in *Med12* hypomorphic mutant mice during development ^56^.

Interestingly, scRNA data from *MED12* mutant UL fibroblasts and smooth muscle cells showed dysregulation of REST targets as we had reported earlier ^9,14^, indicating that PRICKLE1 – REST pathway plays an important role in UL pathogenesis. Despite previous reports on the formation of smooth muscle tumors in mouse model with constitutive activation of β-catenin ^57^, a direct role for WNT/ β-catenin pathway in the pathogenesis of human UL remains unproven. It is conceivable that suppression of DVL1 (Fig S7. C) as well as the overexpression of WNT inhibitory molecules maintain canonical WNT/ β- catenin pathway under control in UL. On the other hand, the role of disrupted WNT/PCP signaling, which may play a crucial role in aberrant tissue architecture and ECM deposition in UL, requires further in-depth investigation.

In the presence of agonists, ERα positively regulates a vast majority of its direct targets in the uterus, contributing to the uterotrophic effects of estrogen. Several selective estrogen receptor modulators (SERMs) have been investigated as potential treatment for uterine fibroids with mixed to moderate success ^58^. The SERMs inhibit the effect of estrogen in the uterus by promoting the recruitment of co-repressors to the estrogen receptor bound transcriptional targets. Since PRICKLE1 seems to be negatively regulated by estradiol and environmental estrogens, it may be crucial to test in the future how PRICKLE1, REST, and the epigenetic status of downstream REST target genes are affected by novel SERMs. Our findings on the role of PRICKLE1 and environmental factors on the stability of REST may have wider implications in hormone dependent cancers and other critical conditions such as Alzheimer’s disease where loss of REST has been reported ^46^. Taken together, our results provide a novel mechanism for the loss of REST and identify potential targets for the development of treatment for fibroids in the future.

## MATERIALS AND METHODS

### Chemicals and reagents

Dulbecco’s Modified Eagle’s medium (DMEM; D5671), penicillin-streptomycin (17-602), and L-glutamine (17-605) were purchased from Biowhittaker (Walkersville, MD). Dulbecco’s PBS (SH30028), fetal bovine serum (FBS; SH30071) and bovine calf serum (BCS; SH30073) were purchased from Hyclone (Logan, UT). Anti-PRICKLE-1 (506-520), anti-rabbit polyclonal antibody (R3782), protease and phosphatase inhibitor cocktail (P0044) and anti-Flag M2 antibody (F1804) were purchased from Sigma-Aldrich (St. Louis, MO). Anti-REST rabbit polyclonal antibody (07-579) and anti-Trimethyl-Histone H3 (Lys27) (07-449) was purchased from Millipore (Temecula, CA). Anti-Actin antibody (sc-47778) and anti-goat IgG HRP conjugated antibody (sc-2020) were purchased from Santa Cruz Biotechnology (Santa Cruz, CA). Ethidium homodimer (E3599), Alexa Fluor conjugated secondary antibodies (ab150073), lipofectamine 2000 reagent (11668027), collagenase type II (17101-015), and high-capacity cDNA reverse transcription kit (4368814) were purchased from Life Technologies (Grand Island, NY). Anti-rabbit IgG HRP conjugate antibody (W401B) was purchased from Promega (Madison, WI). pGateway 3XFlag Prickle1 vector (24644) generated in the lab of Jeff Wrana ^59^ was obtained from Addgene (Cambridge, MA) . TaqMan gene expression master mix (4304437) was purchased from Applied Biosystems (Foster City, CA). NE-PER nuclear extraction reagent (78833) and SuperSignal West Pico Chemiluminescent Substrate (34080) were purchased from Thermo Scientific (Rockford, IL). DR (4084L), *EZH2* (D2C9) XP Rabbit mAb, and *GAPDH (D16H11)* pAb (5174) were purchased from Cell Signaling (Danvers, MA) Taqman primer probe sets, PRICKLE-1 siRNAs (HSC.RNAI.N001144881.12.2, HSC.RNAI.N001144881.12.4, and HSC.RNAI.N001144881.12.8) and *EZH2* siRNAs (HSC.RMAI.N152988.12.2_2nm, HSC.RMAI.N152988.12.7_2nm, HSC.RMAI.N152988.12.8_2nm) were purchased from IDT (Coralville, IA). Gradient 4-15% mini-PROTEAN TGX gels (456-1086) were purchased from Bio-Rad (Hercules, CA). RNeasy Mini Kit (74104) was obtained from Qiagen (Valencia, CA). Antigen unmasking solution (H-3300) was purchased from Vector (Burlingame, CA).

### Tissue collection and cell culture

Leiomyoma tissue samples were obtained from hysterectomies of pre- menopausal women at The University of Kansas Medical Center (Kansas City, KS). Primary smooth muscle tissue cells were prepared from the samples as described previously ^60^.

### Immunoflourescent staining

Human and mice uterus tissue sections were fixed in 4% paraformaldehyde and paraffin embedded. Deparaffinization was with xylene and rehydration with graded alcohol. Samples were then boiled in 1% antigen unmasking buffer.

Tissue sections were blocked in5% normal goat serum and 2% BSA in PBS for 1 hour, then incubated with anti-PRICKLE-1 polyclonal antibody (1:100) or anti-REST polyclonal antibody (1:100) overnight at 4 C. Fluorescence was developed using AlexaFlour 488 conjugated secondary antibody (1:300). Nuclei were stained with is Ethidium Homodimer (1:300) or DR (1:500). An Olympus 1X71 inverted microscope and TE2000U3 laser inverted confocal microscope were used for imaging.

### Protein Isolation and Western Blotting

Leiomyoma and myometrial tissues were homogenized, and primary cells were lysed, in 1x cell lysis buffer containing 1% protease and phosphotase inhibitor cocktails. Supernatant was collected and the pellet was used for nuclear extraction using NE-PER nuclear extraction buffer. Cytoplasmic and nuclear extracts were combined and loaded with 6x Laemmli buffer on 4-15% polyacrylamide SDS gels, run by electrophoresis at 70 volts at room temperature. Transfer to PVDF membrane was at 4°C for 1.5 hours. Membranes were blocked in 5% skim milk for 1 hour followed by an overnight incubation at 4°C with anti-PRICKLE-1 (1:6000), anti-REST (1:2000), anti-FLAG (1:1000), anti-EZH2 (1:1000), anti-H3K27me3 (1:1000), anti-GAPDH (1:1000) or anti-actin (7.5:10000). Secondary antibody incubation was with IgG HRP conjugated antibodies (1: 5000) at room temp for 1 hour. Signal was developed with SuperSignal West Pico Chemiluminescent Substrate. ImageJ software from the National Institutes of Health was used for densitometric analysis.

### Cell transfections

Primary fibroid or myometrial cells were cultured to 70% confluency in 6-well plates and incubated overnight in antibiotic-free DMEM medium before transcfection with 25nM *PRICKLE-1* siRNA (HSC.RNAI.N001144881.12.2, HSC.RNAI.N001144881.12.4, and HSC.RNAI.N001144881.12.8; IDT) or *EZH2* siRNA (HSC.RMAI.N152988.12.2_2nm, HSC.RMAI.N152988.12.7_2nm, HSC.RMAI.N152988.12.8_2nm) or 4ug of pGateway 3XFlag Prickle1 vector using Lipofectamine 2000. SiRNA or vector containing medium was removed 24 hours after transfection and cells were harvested 24 hours later.

### RNA isolation and real time qPCR

RNA was extracted using Qiagen RNeasy Kit. cDNA was prepared with the High Capacity cDNA Reverse Transcription Kit using 1μg RNA. Probes for total *PRICKLE-1* (Hs.PT.49a.20111671)*, PRICKLE-1* mRNA 3’UTR variants (nm153026.1.pt.prickle1, nm001144882.1.pt.prickle1), *REST* (Hs.PT.47.3525906: IDT), and *EZH2* (Hs.PT.58.1924301) were used in combination with Taqman gene expression master mix. Four experimental replicates were performed for each sample. An ABI 7900HT sequence detection system was used for real time PCR amplification. The comparative CT method (ΔΔCt) was applied for data analysis. Relative fold differences in gene expressions were normalized to 18S (Hs.PT.49a.3175696.g; IDT) expression as an internal control.

### Animals

Animals were handled according to National Institutes of Health National Institute of Environmental Health Sciences guidelines under approved animal care and use protocols. Female CD-1 mice were injected subcutaneously on PND1–PND5 with corn oil (control) or Genistein (50 mg/kg/day; Sigma, St. Louis, MO) or DES (1 mg/kg/day; Sigma). Control and Gen-treated uteri were collected at 5 days of age and fixed in 4% paraformaldehyde ^61^. Tissues were embedded, sectioned and processed as described in the histology section. Additional control and DES-treated mice were aged to 8 weeks of age, ovariectomized to remove endogenous hormones, and 10-14 days later were treated with vehicle (corn oil) or estradiol (Sigma) at a dose of 25 µg/mouse for 24 hours. Uteri were collected and frozen at -80C. RNA was isolated as described RNA isolation section.

*Esr1-/-* mice (B6.129P2-Esr1tm1Ksk/J) were purchased from The Jackson Laboratory. Uteri of 3-4 month old female *Esr1-/-* mice were isolated and processed for histology as described above.

*Lhb-/-* mice were generated and genotyped as described previously ^62^. Adult female Lhb-/- or littermate control mice were treated with 5μg/kg estradiol or placebo time release pellets for 7 days before sacking and uteri collection.

### EZH2, H3K27Me3 ChIP-PCR

Chromatin immunoprecipitation experiments were performed in matched myometrial and UL tissue samples as described previously ^9,14^. The distal TSS sequences of *PRICKLE1* overlapping ERα binding regions were amplified using primers: FWD primer CCCGTTTGGACTCGATCCTG and REV primer GGGAACGGACTAGAGGCTG (which amplify 119 base pairs of human chromosome 12:42984112- 42984231); and FWD primer TTCCGGATCGGTGGAGTCTC and REV primer AGAGGCACCTGTCAAGTTCG (which amplify 100 base pairs of human chromosome 12:42984550- 42984650). Additionally, the proximal TSS sequences of *PRICKLE1* sequences were amplified using: FWD primer CTTTCCCTGCATTTCGGAGTG and REV primer AGTTAGTACATCGCGTGACCTG (which amplify 105 base pairs of human chromosome 12: 42878325 – 42878429) and FWD primer GCATAGAACAGTCGCCTTCG and REV primer TTGGAAGCTGAGGGACAGAC (which amplify 160 base pairs of human chromosome 12:42876920- 42877080)

## ACKOWLEDGEMENTS

We thank Stan Fernald for illustrations; and Dr. Jeff Wrana for deposition of 24644 pGateway 3x Flag Prickle1 plasmid in the Addgene plasmid repository. VMC was supported by grants from the NIH: P20 RR016475, R01 HD094373, R01HD076450. Authors acknowledge University of Kansas (KU) Cancer Center Biospecimen Repository Core for human specimens, KU Cancer Center’s Support Grant (P30 CA168524), Genomics Core supported by the Kansas Intellectual and Developmental Disability Research Center (NIH U54 HD090216), COBRE (P30 GM122731-03) and the NIH S10 High-End Instrumentation Grant (NIH S10 OD021743) at KUMC, Kansas City, KS 66160. MMM was supported by The Madison and Lila Self Graduate Fellowship.

---

Fig. S1. PRICKLE-1 mRNA expression is decreased in uterine leiomyomas. Gene expression analysis of PRICKLE1 in myometrium and leiomyoma tissue samples (12 pairs). * P < 0.05

Fig. S2. REST localization is primarily cytoplasmic in leiomyomas. (A & B) Immunofluorescence analysis of REST (green) in myometrial and leiomyoma tissue samples. (C &D). Immunofluorescence analysis of GFP-REST(green) transfected MSMCs and LSMCs. Nuclei were stained with EthD-2 (red).

Fig. S3. PRICKLE1 regulates REST at the protein level. Gene expression analysis of PRICKLE1 and REST in LSMCs overexpressing FLAG-PRICKLE-1.

Fig. S4. Neonatal (PND1-5) DES exposure suppresses Prickle1 expression in the mouse uterus. Relative mRNA levels from Affymetrics gene expression arrays

Fig. S5. Neonatal exposure to DES augments the suppression of Prickle1 by 17β − estradiol. TaqMan qRT-PCR assay showing relative Prickle1 mRNA expression in mice treated neonatally (PND1-5) with vehicle or DES, ovariectomized at 8 weeks of age and then treated with vehicle or 17β - estradiol. Effect of estradiol on Prickle1 mRNA expression in control mice treated neonatally (PND1-5) with vehicle (samples; veh, E2), effect of estradiol on Prickle1 mRNA expression in adult ovariectomized mice which were treated neonatally (PND1-5) with DES (samples; DES, DES + E2). Dotted lines indicate P< 0.05

Fig. S6. EZH2 mediated repression of PRICKLE1 in leiomyomas. A. UCSC genome browser data showing H3K27Me3 peaks near the major TSS sequences of PRICKLE1. B, C, representative ChIP PCR from patients showing association of EZH2 to the distal start site (red box in A) and increased H3K27 methylation in UL.

Fig. S7. Disruption of WNT/ planar cell polarity pathway in leiomyomas. A. Immunofluorescence staining of β-catenin showing predominant cytoplasmic localization in UL tissue. B. Analysis of gene expression dataset GSE13319 showing downregulation (green) or upregulation (red) of TCF/LEF targets. C. Western blot analysis showing the absence of LRP5/6 activation and the suppression of disheveled 1 in UL. D. TaqMan qRT-PCR data showing expression of TCF/LEF targets. E, F downregulation of DVL1 mRNA and protein respectively in UL. G, H. expression of WNT inhibitory molecules and WNT/ PCP targets in UL respectively. I. Western blot showing regulation of WNT/PCP signaling in UL.

