## Supplementary figures and images for "Estrogen receptor alpha mediated repression of PRICKLE1 destabilizes REST and promotes uterine fibroid pathogenesis"

### Supplemental data

**Supplemental Figures**

Fig. S1

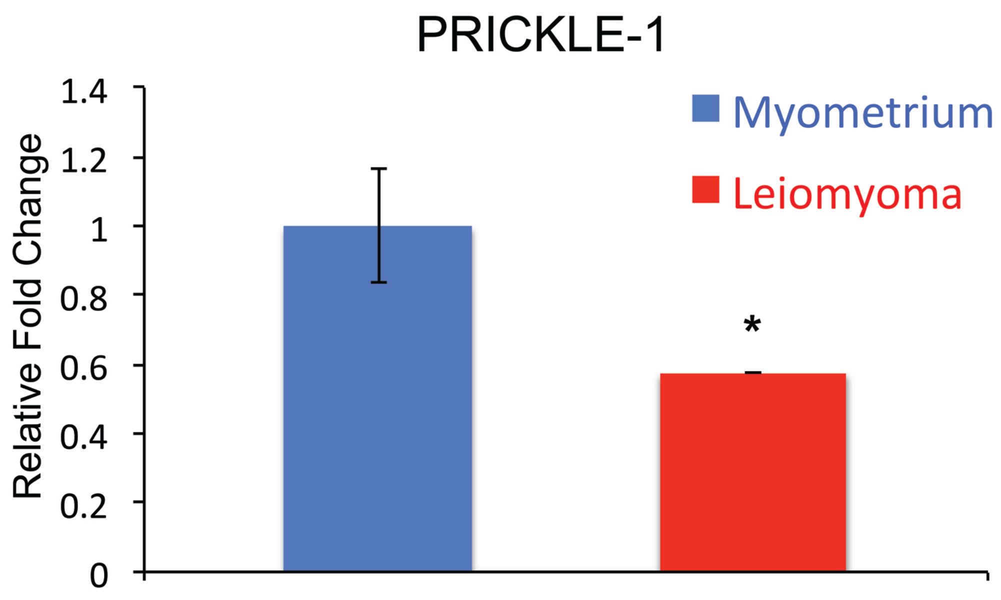

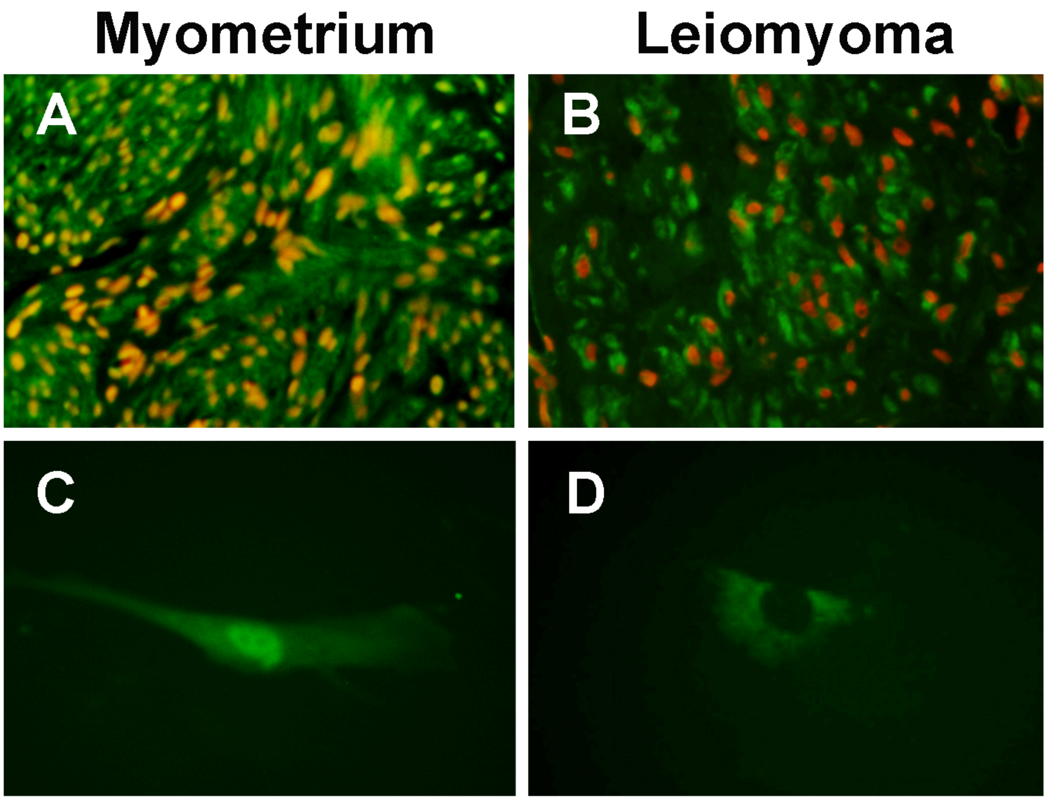


Fig. S2

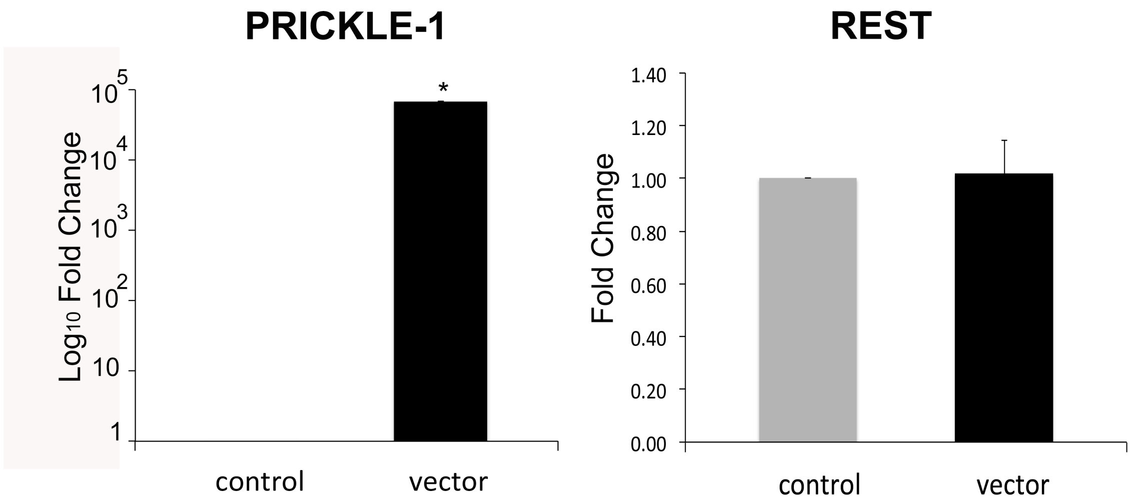

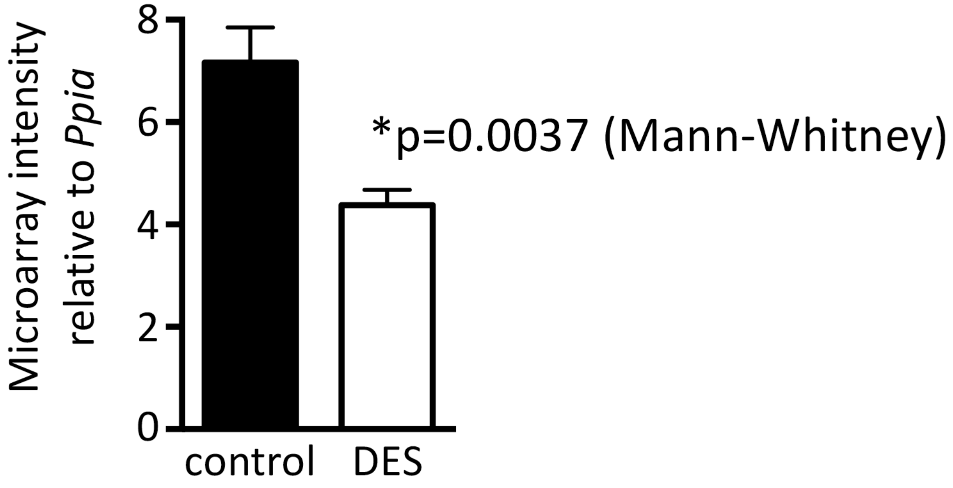

Fig. S5

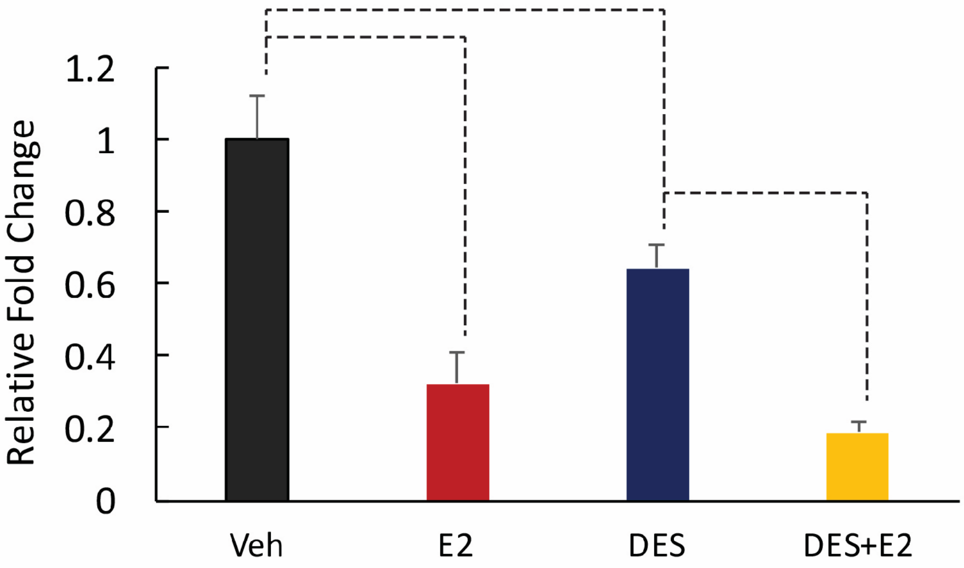

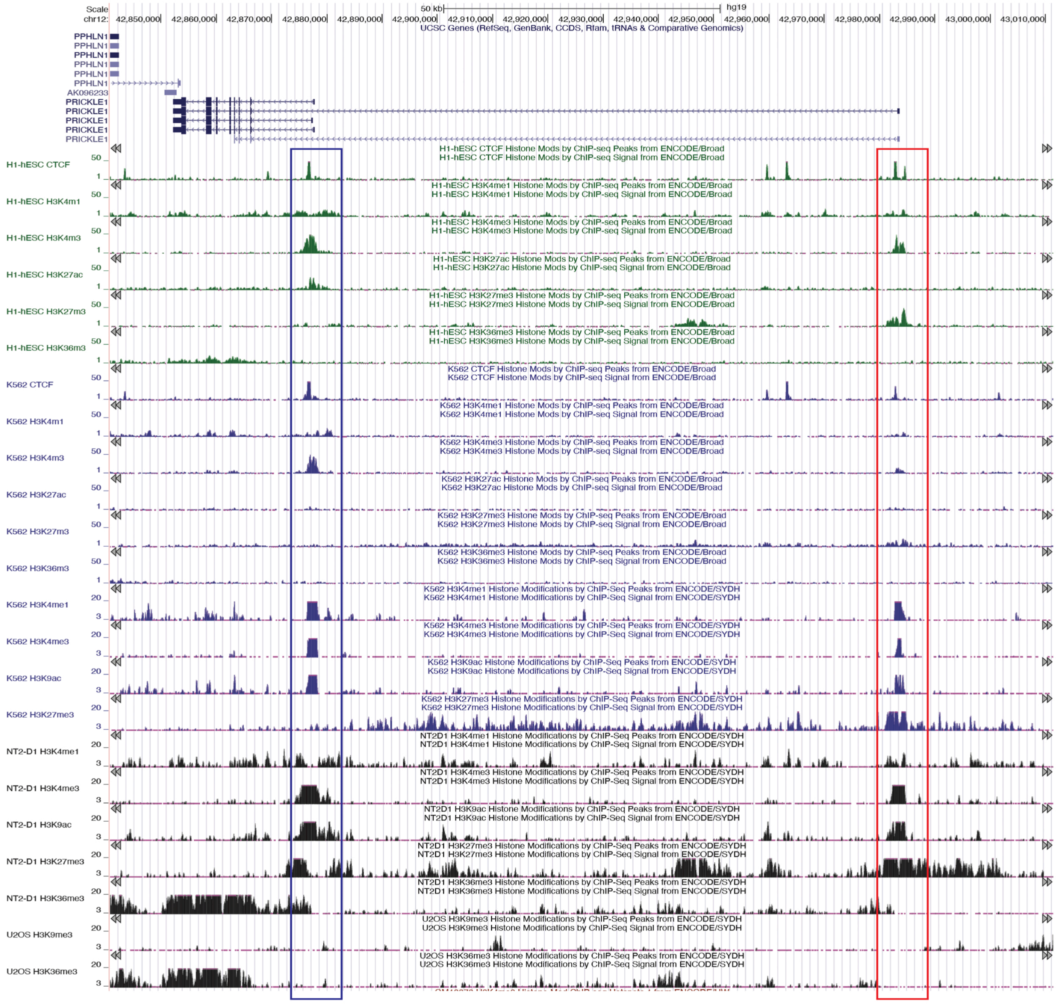

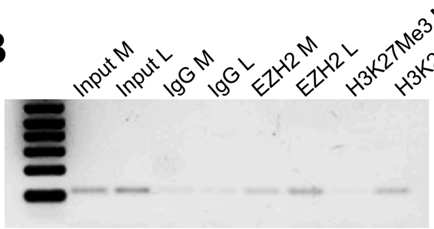

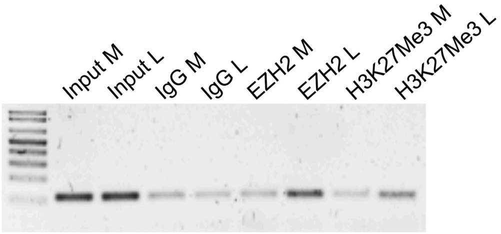

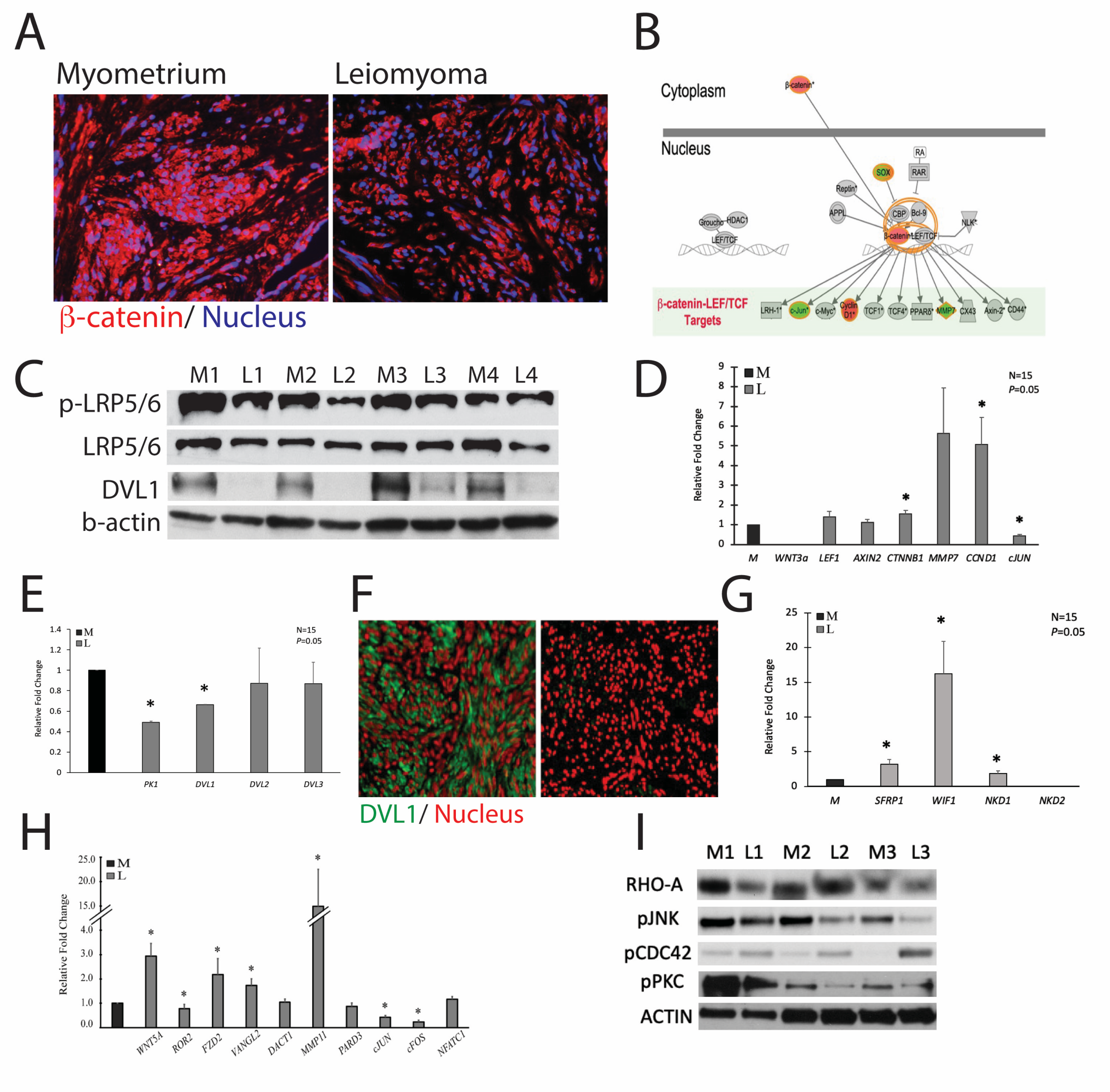
